# Selective vulnerability of excitatory cortical neurons to ZMYND11 loss reveals cell-type-specific mechanisms of neurodevelopmental disorder risk

**DOI:** 10.64898/2026.06.16.732737

**Authors:** Xuyao Chang, Megan McKinley, Wenqi Li, Inah Yang, Elena Albizzati, Eri Iwasawa, Feng Guo, Amelle Shillington, Jason Tchieu

## Abstract

Excitation/inhibition (E:I) imbalance is a convergent mechanism in neurodevelopmental disorders (NDDs), yet whether NDD risk genes disrupt excitatory and inhibitory neurons through shared or distinct molecular programs remains poorly understood. Using human pluripotent stem cell-derived cortical projection and medial ganglionic eminence-like inhibitory neurons, we show that loss-of-function mutations in the chromatin reader ZMYND11 produce cell-type-selective vulnerability in cortical excitatory neurons. ZMYND11-deficient excitatory neurons exhibit hyperexcitability accompanied by de-repression of BMP signaling, dysregulation of glutamate receptor expression, and a shift toward non-brain splicing isoforms, whereas these molecular signatures are largely absent in ZMYND11-deficient inhibitory neurons. This cell-type selectivity is associated with differential upregulation of RBFOX family splicing regulators.

Structure-function analysis reveals that MYND domain is required for normal progenitor dynamics and neuronal excitability. Together, these findings indicate that E:I imbalance in ZMYND11-associated NDD arises primarily from cell-type-selective vulnerability of excitatory cortical projection neurons and identify MYND domain as a critical determinant of neuronal function.

**Highlights:**

- ZMYND11-deficient excitatory neurons (ExNs) show hyperexcitability.
- Altered BMP signaling & glutamate receptor levels are implicated in mutant ExNs.
- Inhibitory neurons show attenuated response to ZMYND11 loss.
- The MYND domain in ZMYND11 is a critical regulator of neuronal excitability.

## Introduction

Neurodevelopmental disorders (NDDs), including autism spectrum disorder and epilepsy, arise from diverse genetic and environmental etiologies yet often converge on shared circuit-level dysfunction (Bozzi et al., 2018; Sylvester et al., 2025; Tatti et al., 2017; Cline, 2005; Sohal and Rubenstein, 2019; Froemke, 2015). A prevailing model proposes that disruption of the balance between excitatory and inhibitory neuronal activity within the cerebral cortex, commonly referred to as excitation/inhibition (E:I) imbalance, is a core pathophysiological mechanism underlying many of these conditions (Rubenstein and Merzenich, 2003; Uzunova et al., 2016; Antoine et al., 2019; Gatto and Broadie, 2010). Evidence from both clinical studies and preclinical models indicates that many genetic risk factors for NDDs, including single-gene syndromes and copy number variants, alter synaptic development, neurotransmission, or intrinsic neuronal excitability in ways that shift this balance (Pietropaolo and Provenzano, 2022). However, because E:I balance reflects the integrated properties of both excitatory and inhibitory neuronal populations, it remains unclear whether NDD-associated mutations disrupt these lineages through shared molecular mechanisms or through cell-type–specific vulnerabilities. Resolving how genetic perturbations differentially affect excitatory and inhibitory neurons is therefore essential for understanding how diverse risk genes converge to produce circuit-level dysfunction.

Recent human stem cell studies have begun to reveal how NDD-associated mutations influence neuronal excitability (Cheffer et al., 2020; Hussein et al., 2023; Deneault et al., 2019; Lamar and Carvill, 2018; Sarkar et al., 2022), but they also highlight an important gap in understanding how these effects differ across neuronal lineages. Early iPSC-based models of autism reported increased spontaneous activity in cortical neurons and forebrain organoids, with purified excitatory neurons exhibiting particularly robust hyperexcitability and network bursting compared with mixed neuronal populations (Shcheglovitov et al., 2013; Yi et al., 2016; Fink et al., 2021; Santos et al., 2023; Avazzadeh et al., 2021). Larger-scale analyses of ASD-associated mutations similarly identified network hyperactivity in iPSC-derived excitatory neurons but emphasized the lack of parallel inhibitory interneuron models from the same genetic backgrounds (Deneault et al., 2019; Culotta and Penzes, 2020; Nelson and Valakh, 2015). More recent work has begun to address this issue by deriving excitatory and inhibitory neurons from the same patient lines, demonstrating that mutations in epilepsy-associated genes such as *SCN1A* can produce divergent cell-type–specific excitability defects, hyperexcitability in excitatory neurons and reduced firing in inhibitory neurons, resulting in overall network imbalance (Sun et al., 2016; Kim et al., 2018; Jiao et al., 2013). Consistent with these findings, large-scale genetic and transcriptomic analyses of autism risk genes indicate enrichment across both excitatory and inhibitory neuronal populations and suggest that differential cell-type vulnerability may contribute to disease mechanisms (Gandal et al., 2022; Wang et al., 2018; Satterstrom et al., 2020). Notably, many high-confidence NDD risk genes encode chromatin regulators expressed early in neural progenitors (Cederquist et al., 2020), raising the possibility that shared developmental perturbations may produce lineage-specific consequences in downstream neuronal populations. In support of this concept, our recent work showed that loss of the chromatin reader ZMYND11 disrupts progenitor proliferation and neuronal differentiation through dysregulation of BMP signaling and RNA splicing programs in human neural models (Chang et al., 2025). However, whether ZMYND11 mutations produce equivalent or divergent functional consequences in excitatory versus inhibitory neuronal lineages remains unknown.

Here, we address this question by deriving excitatory and inhibitory neurons from isogenic *ZMYND11* mutant human pluripotent stem cells (hPSCs) and systematically comparing their functional and molecular responses to ZMYND11 loss. Using calcium imaging, bulk RNA sequencing, and alternative splicing analysis, we find that ZMYND11 deficiency produces hyperexcitability selectively in excitatory neurons. This phenotype is accompanied by de-repression of developmental signaling pathways, dysregulation of glutamate receptor gene expression, and a shift toward non-brain splicing isoforms. In contrast, inhibitory neurons exhibit a markedly attenuated molecular response, with no enrichment of developmental signaling pathways and minimal overlap in differentially expressed genes. Structure-function analysis further reveals that the C-terminal MYND domain, a hotspot for patient missense mutations, is required for normal neuronal excitability and loss of this domain recapitulates the transcriptomic profile of heterozygous patient-like mutations. Together, these findings establish cell-type specific vulnerability of excitatory neurons as a source for E:I imbalance in ZMYND11-associated NDDs.

## Results

### ZMYND11 deficiency induces hyperexcitability in human excitatory neurons

We previously showed that ZMYND11 is expressed throughout human corticogenesis and mutations impair the generation of secondary progenitors (Chang et al., 2025). ZMYND11-deficient neural progenitors display aberrant BMP signaling and inhibition of this pathway rescues the neurogenesis deficit, enabling efficient derivation of excitatory neurons from dorsal neural progenitors. To determine whether ZMYND11 mutations affect neuronal function beyond the progenitor stage, we used this established protocol to generate cortical excitatory neurons from isogenic wild-type, heterozygous (ZMYND11 +/−), and knockout (ZMYND11 −/−) hESC lines as described before (Chang et al., 2025). Cortical neural progenitors were patterned toward a prefrontal identity using dual-SMAD inhibition with WNT antagonism followed by FGF8 regionalization until day 18, then driven to terminal differentiation by NOTCH inhibition using DAPT for 5 days (**Figure 1A**). The resulting neurons expressed the deep-layer cortical marker TBR1 and the pan-neuronal marker MAP2, confirming appropriate regional and neuronal identity (**Figure 1B**). Importantly, after further maturation till day 60, ZMYND11-deficient excitatory neurons showed comparable expression of SYN1 relative to controls (**Figure S1A**), indicating that maturation state was not grossly altered by ZMYND11 loss and that functional differences would not be attributable to delayed maturation.

**Figure 1.**
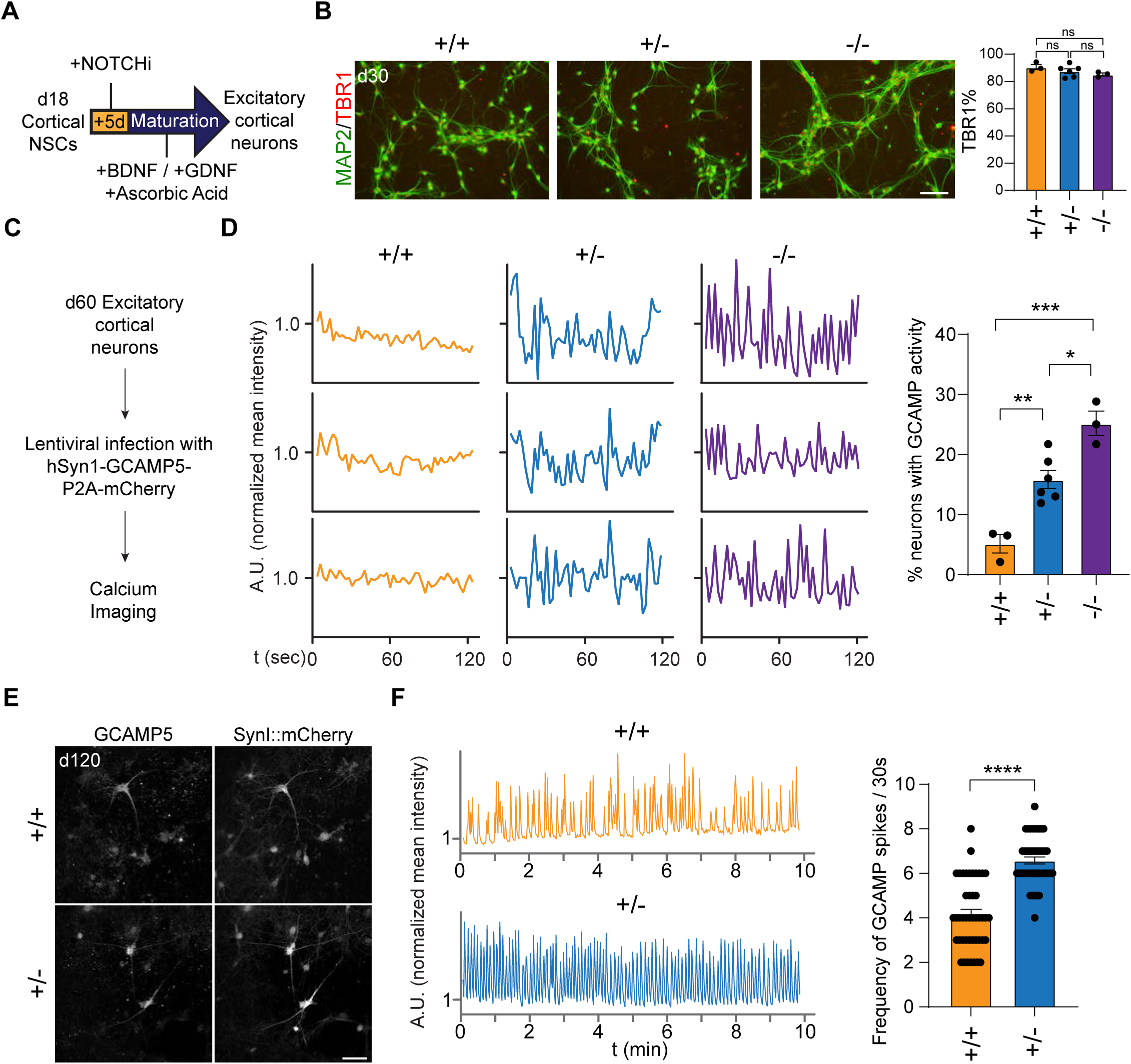
ZMYND11 deficient excitatory neurons show hyperexcitability. (**A**) Schematic outline of excitatory neuron induction protocol from cortical neural stem cells (NSCs) at day 18 (d18). (**B**) Immunostaining of MAP2 (green) and TBR1 (red) on excitatory neurons at day 30 (d30) with quantification (mean ± SEM, n= 3 for +/+ and -/-, n= 6 for +/-). Scale bar, 50μm. (**C**) Schematic outline of hSYN1-GCAMP5-P2A-mCherry virus infection and calcium imaging at day 60 (d60). (**D**) Representative normalized traces of spontaneous calcium handling at ∼d60 with quantification of the number of firing neurons (mean ± SEM, n= 3 for +/+ and -/-, n= 6 for +/-). (**E**) Representative imaging of GCAMP5 and mCherry on day 120 (d120) neurons. Scale bar, 50μm. (**F**) Representative normalized traces of spontaneous calcium handling at d120 with quantification of frequency of GCAMP spike per 30 seconds (mean ± SEM, n= 45 for +/+ and +/-). Each n in **(B)** and **(D)** represents an independent differentiation, and in **(F)** represents an individual neuron across three independent differentiations. *P* values in **(B)** and **(D)** were calculated by one-way ANOVA followed by Tukey’s test, while in **(F)** by unpaired two-tailed t test (two groups). ns: not significant. \**P*< 0.05; \*\**P*< 0.01; \*\*\**P*< 0.001; \*\*\*\**P*< 0.0001. See also **Figure S1**.

Having confirmed that ZMYND11-deficient excitatory neurons differentiated and matured comparably to controls, we next asked whether they exhibit altered functional properties. To assess neuronal activity, we transduced neurons with a lentiviral calcium reporter (hSYN1-GCaMP5-P2A-mCherry) and performed live calcium imaging of spontaneous activity (**Figure 1C**). The mCherry reporter served as both an infection marker and a proxy for synapsin-1-driven expression, ensuring that imaged neurons had reached a baseline level of synaptic maturity. At approximately day 60 of differentiation, ZMYND11-deficient excitatory neurons displayed a significantly greater proportion of spontaneously active cells compared with wild-type controls, further increased with ZMYND11 loss, and these active neurons exhibited increased calcium transient amplitude, indicative of increased spontaneous calcium activity (**Figure 1D and Videos S1-S3**). To determine whether hyperexcitability persisted or resolved with further maturation, we continued culturing neurons to approximately day 120 (**Figure 1E**). At this later point, ZMYND11-deficient excitatory neurons continued to exhibit significantly elevated spontaneous calcium activity compared with controls (**Figure 1F and Videos S4-S5**), indicating that the hyperactivity phenotype is not a transient feature of early neuronal differentiation but persists during neuronal maturation. The persistence of this phenotype across a 60-day maturation window suggests that ZMYND11 deficiency produces a persistent hyperactive neuronal state.

### ZMYND11 loss activates developmental signaling and alters glutamatergic molecular programs in excitatory neurons

To identify molecular programs underlying this hyperexcitability, we performed bulk RNA sequencing on ZMYND11-deficient and control excitatory neurons at day 60. As expected, *SYN1* expression was similar between control and mutants (**Figure S1B**). Principal component analysis revealed clear transcriptomic separation between mutant and control samples (**Figure 2A**). We identified 152 upregulated and 132 downregulated genes in heterozygous mutants and 493 upregulated and 294 downregulated genes in knockout neurons (**Figure 2B and Table S1**). Although fewer genes were dysregulated than in ZMYND11-deficient cortical neural progenitors in our previous study, genes that remain altered in postmitotic neurons likely represent particularly robust targets of ZMYND11 regulation. Consistent with this idea, a substantial fraction of differentially expressed genes was shared between heterozygous and knockout neurons, suggesting dose-sensitive regulation of a core gene set (**Figure 2C**).

**Figure 2.**
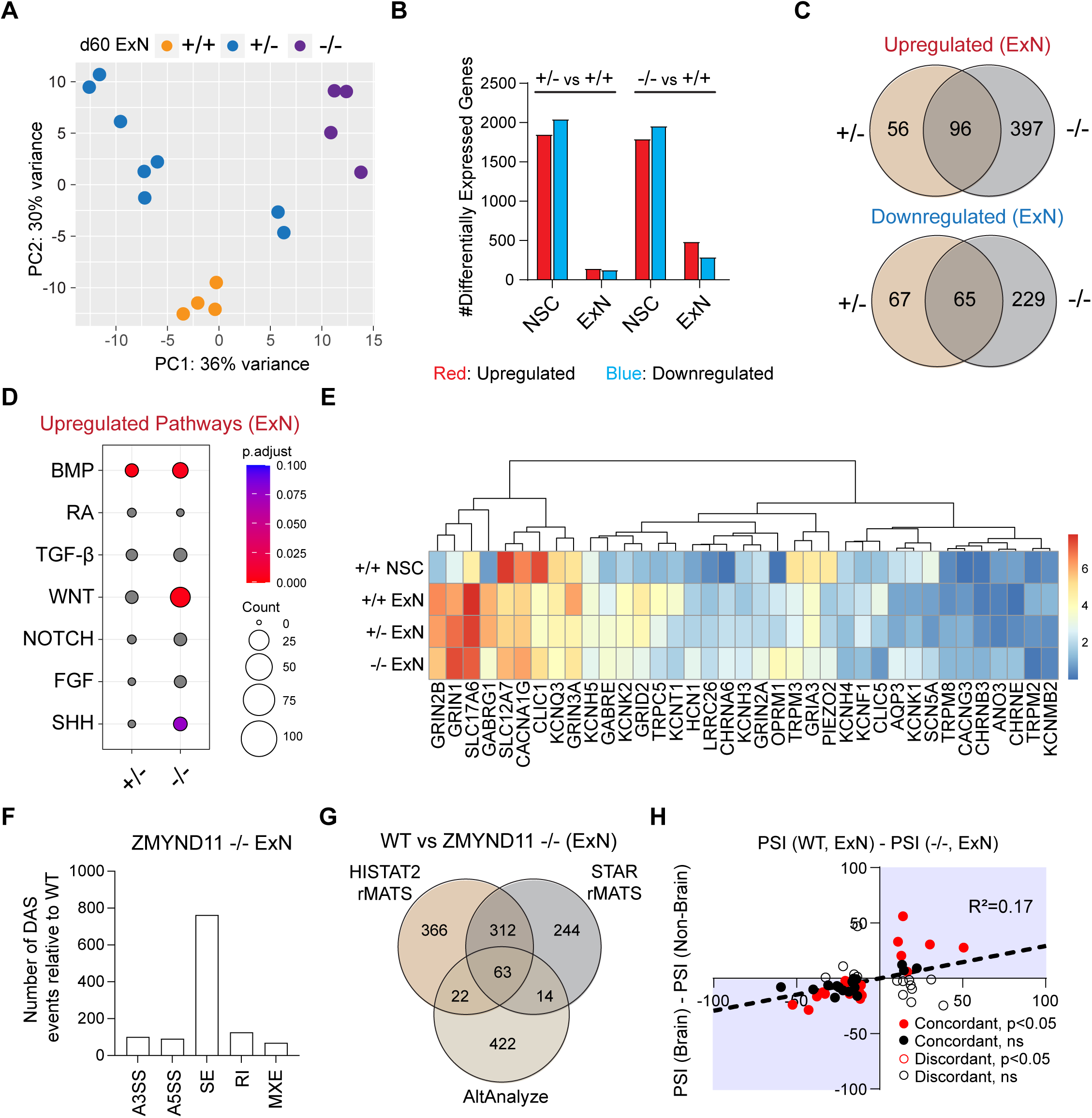
Transcriptomic analysis on ZMYND11 deficient excitatory neurons reveals latent developmental signaling activation, alteration of channel protein expression and non-brain isoform switch. **(A)** PCA of RNA-seq on control and mutant excitatory neurons at day 60 (d60, n= 4 for +/+ and - /-, n= 8 for +/-). **(B)** Quantification of the number of differentially expressed genes in cortical neural stem cells (NSCs) and excitatory neurons. **(C)** Overlap of differentially expressed genes in ZMYND11 deficient excitatory neurons (*P* value < 1e-80). **(D)** Gene ontology analysis on upregulated genes in ZMYND11 deficient excitatory neurons. Gray dots: *P*.adjust > 0.1. **(E)** Heatmap of ion channel protein expression (log2(cpm+1)) across mutants and control. **(F)** Number of differential alternative splicing (DAS) events in ZMYND11 -/- excitatory neurons (rMATS-HISAT2). A3SS, A5SS: Alternative 3’/5’ splice site. SE: Spliced exon. RI: Retained intron. MXE: Mutually exclusive exons. **(G)** Overlap of DAS events predicted by HISAT2-rMATS, STAR-rMATS and AltAnalyze in ZMYND11 -/-. **(H)** Scatter plot of PSI differences between WT vs. KO excitatory neurons and brain vs. non-brain tissues. Red/black marks significant (P < 0.05) or non-significant events, respectively. Closed/open circles show matched vs. unmatched KO and non-brain tissue trends. Each n represents an independent differentiation. See also **Figure S2**.

Because ZMYND11 functions as a transcriptional co-repressor, we focused our gene ontology (GO) analysis on upregulated genes to identify programs that are normally repressed by ZMYND11. Consistent with our findings in cortical neural progenitors, we observed significant enrichment of developmental signaling pathways, including BMP and WNT in ZMYND11-deficient lines compared to controls (**Figures 2D and S2A**). De-repression of these pathways in postmitotic neurons is notable because BMP and WNT signaling can modulate synaptic transmission and neuronal excitability in mature circuits (Okur et al., 2024; Karwacki-Neisius et al., 2024), raising the possibility that persistent developmental signaling contributes directly to the hyperexcitability phenotype. In addition, GO analysis revealed enrichment of non-neuronal developmental terms among upregulated genes (**Figure S2B**), consistent with a broader loss of transcriptional repression of non-neuronal programs.

We next examined whether changes in ion channel gene expression could provide a more direct link to the excitability phenotype. Among differentially expressed genes enriched in the gene ontology term "channel activity," we identified discordant expression changes across multiple glutamate receptor subunits (**Figure 2E**). *GRIN1* and *GRIN2A* were upregulated in ZMYND11 mutants, while *SLC17A6*, *GRIN3A*, *GRID2*, *GRIN2B*, and *GRIA3* were downregulated. This pattern is potentially significant for several reasons. *GRIN1* and *GRIN2A* encode obligatory and regulatory NMDA receptor subunits, and their co-upregulation could increase NMDA receptor-mediated currents and calcium influx (Dupuis et al., 2023). In contrast, downregulation of the inhibitory subunit *GRIN3A*, which attenuates NMDA receptor conductance, may further potentiate excitatory signaling. Notably, *GRIN1*, *GRIN2A*, *GRID2*, *GRIN2B*, and *GRIA3* are all cataloged in the SFARI database as autism risk genes (Abrahams et al., 2013), highlighting dysregulation of glutamatergic signaling as a convergent mechanism in NDD pathogenesis.

Beyond gene expression changes, we asked whether ZMYND11 loss disrupts alternative splicing in excitatory neurons. Using a stringent pipeline combining rMATS (Shen et al., 2014) and AltAnalyze (Emig et al., 2010) to identify high-confidence differential alternative splicing (DAS) events, we detected 63 events in knockout neurons and 23 in heterozygous mutants (**Figures 2F-2G**, **S2C-S2D and Table S2**). This represents a substantial reduction compared to the 590 DAS events observed in ZMYND11-deficient neural progenitors (Chang et al., 2025), suggesting that the alternative splicing phenotype is attenuated but not lost in ZMYND11-deficient postmitotic neurons. To characterize the directionality of these splicing changes, we correlated the percent-spliced-in (PSI) values of DAS events in ZMYND11 mutants with tissue-specific PSI values from the *VastDB* database (Tapial et al., 2017). In knockout excitatory neurons, we observed a significant shift toward non-brain splicing isoforms (**Figure 2H**), recapitulating the non-brain isoform switch we previously identified in ZMYND11-deficient cortical NSCs (Chang et al., 2025). This shift was not observed in heterozygous mutant excitatory neurons (**Figure S2E**), suggesting dose-dependent severity. Genes harboring high-confidence DAS events in excitatory neurons were enriched in cytoskeletal and cilium-related functions (**Figure S2F**) and did not overlap with DAS genes identified in cortical NSCs **(Figure S2G)**, indicating that while the overall directionality of the splicing phenotype is conserved across developmental stages, the specific gene targets are cell-type dependent.

Together these analyses show that ZMYND11-deficient excitatory neurons exhibit de-repression of developmental signaling pathways, altered glutamate receptor expression, and a shift toward non-brain isoforms. Whether these changes represent a general neuronal response to ZMYND11 loss or a lineage-specific vulnerability remain unclear.

### Inhibitory neurons exhibit an attenuated and divergent response to ZMYND11 loss

To directly test whether the molecular and functional consequences of ZMYND11 loss are shared across neuronal lineages or specific to excitatory cortical neurons, we generated inhibitory neurons from the same isogenic ZMYND11 mutant lines. Ventral progenitors were generated by adding the sonic hedgehog agonist SAG during cortical differentiation from day 8 to 18, yielding FOXG1 positive, NKX2-1 positive neural progenitors that lacked expression of the dorsal marker PAX6 (**Figures 3A and S3A**) and thus, likely represented medial ganglionic eminence (MGE)-like cortical interneuron progenitors. Upon NOTCH inhibition for 5 days, these progenitors differentiated into GAD67-expressing inhibitory neurons, in contrast to the VGLUT2-expressing excitatory neurons produced from dorsal progenitors (**Figure 3B**).

**Figure 3.**
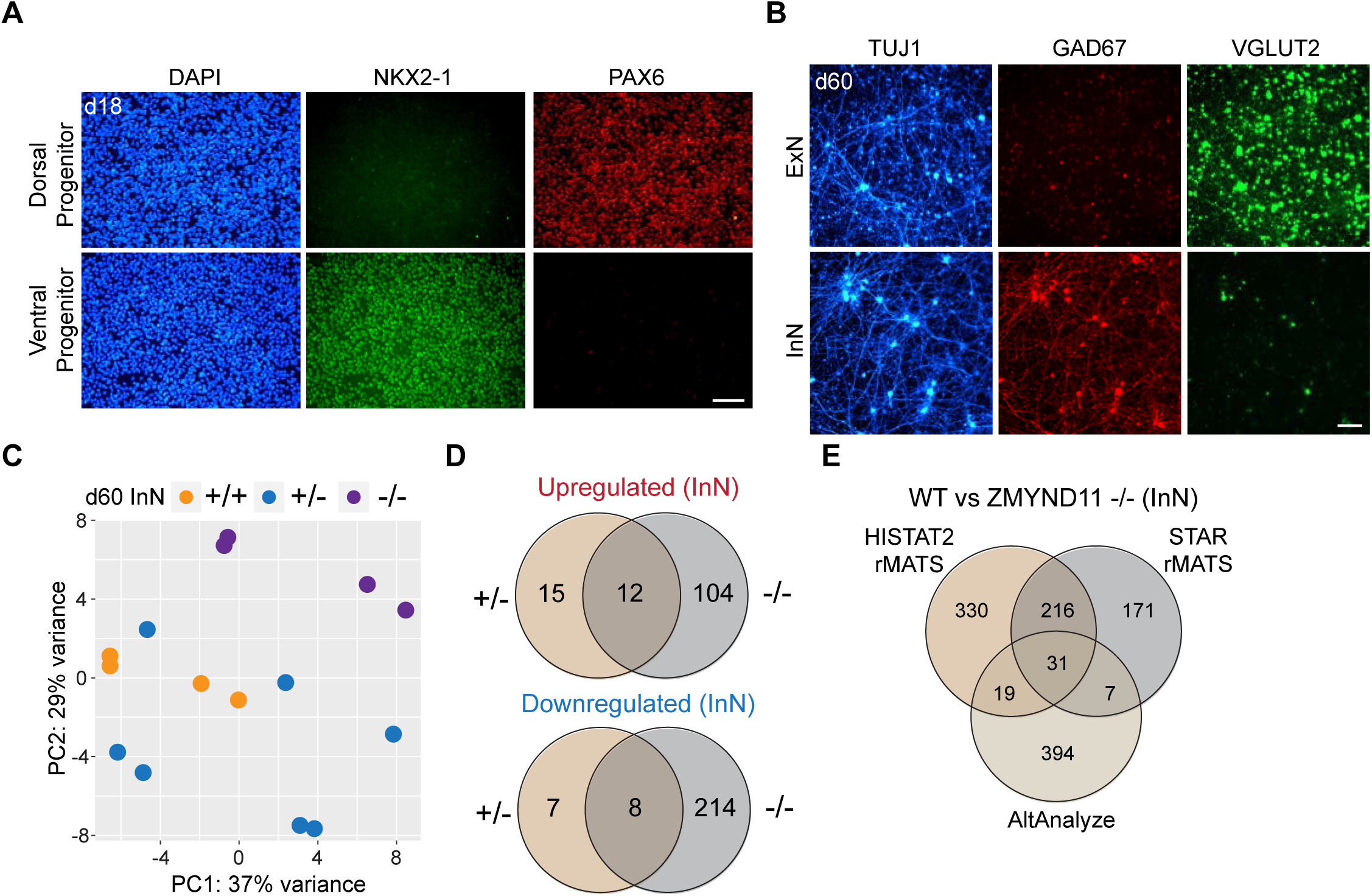
Transcriptomic analysis on ZMYND11 deficient inhibitory neurons reveals less susceptibility to the loss of ZMYND11 compared with excitatory neurons. (**A**) Immunostaining of NKX2-1 (green) and PAX6 (red) on wildtype (WT) dorsal progenitors and ventral progenitors at day 18 (d18, n=3). Scale bar, 50μm. (**B**) Immunostaining of TUJ1 (blue), GAD67 (red) and VGLUT2 (green) on WT excitatory neurons and inhibitory neurons at day 60 (d60, n=3). Scale bar, 100μm. (**C**) PCA of RNA-seq on control and mutant inhibitory neurons at day 60 (d60, n= 4 for +/+ and - /-, n= 7 for +/-). (**D**) Overlap of differentially expressed genes in ZMYND11 deficient inhibitory neurons (P value < 1e-10). (**E**) Overlap of DAS events predicted by HISAT2-rMATS, STAR-rMATS and AltAnalyze in ZMYND11 -/- inhibitory neurons. Each n represents an independent differentiation. See also **Figure S3**.

We performed bulk RNA-sequencing on mutant and control inhibitory neurons at day 60 and found a strikingly different transcriptomic landscape compared with excitatory neurons. Principal component analysis revealed clear separation between knockout and control inhibitory neurons; however, heterozygous mutants were largely indistinguishable from controls (**Figure 3C**). Consistent with this, differential gene expression analysis identified only 27 upregulated and 15 downregulated genes in heterozygous mutants, compared with 116 upregulated and 222 downregulated genes in knockout inhibitory neurons (**Figure 3D and Table S1**). Beyond this reduction, the inhibitory neuron response was qualitatively distinct from that observed in excitatory neurons. GO analysis of upregulated genes revealed no enrichment of BMP, WNT, or other developmental signaling pathways (**Figure S3B**), in contrast to the prominent developmental signaling signature observed in ZMYND11-deficient excitatory neurons. This was notable given that *ZMYND11* expression was higher in inhibitory neurons (**Figure S3C**). Moreover, differentially expressed genes in inhibitory neurons showed minimal overlap with those in excitatory neurons (**Figure S3D**), indicating that ZMYND11 loss perturbs largely non-overlapping gene sets between these two lineages rather than producing a shared transcriptomic response.

Despite the minimal expression changes, we wondered if splicing was disrupted. We performed a similar alternative splicing analysis that reinforced this cell-type distinction. High-confidence DAS events were fewer in inhibitory neurons (31 in knockout, 12 in heterozygous mutants; **Figures 3E**, **S3E and Table S2**), and the directionality of splicing changes was unexpectedly reversed. Whereas ZMYND11-deficient excitatory neurons shifted toward non-brain isoforms, knockout inhibitory neurons showed a trend toward brain-specific isoforms (**Figures S3F-S3G**). To investigate the basis for this divergence, we examined expression of the RBFOX family of RNA-binding proteins, which was shown to regulate brain-specific isoform switch in our previous study (Chang et al., 2025). ZMYND11-deficient excitatory neurons showed significant upregulation of *RBFOX1* and *RBFOX3*, while inhibitory neurons showed no changes in *RBFOX1*, *RBFOX2*, or *RBFOX3* expression (**Figure S3H**). This differential RBFOX dysregulation provides a potential explanation for the opposing splicing trajectories observed in the two cell types. We also examined SRRM4 and its paralog SRRM3, which have been implicated in neuronal isoform switching(Irimia et al., 2014), but detected no expression changes in either excitatory or inhibitory neurons (**Figure S3I**), suggesting that the splicing phenotype operates independently of this pathway.

### MYND domain loss drives hyperexcitability in ZMYND11 deficient excitatory neurons

Our analyses thus far examined the consequences of ZMYND11 haploinsufficiency and complete loss of function. However, many patient variants in *ZMYND11* are missense substitutions rather than truncating mutations, and several cluster within the C-terminal MYND domain (residues 563–598; **Figure S4A**). This enrichment suggests that the MYND domain plays an important role in ZMYND11 function during neurodevelopment, yet its contribution to neuronal phenotypes has not been directly tested. To investigate the structure-function relationship between the MYND domain and the neuronal phenotypes we identified, we used CRISPR/Cas9 to introduce a premature stop codon at the start of the MYND-encoding exon in hPSCs, generating homozygous MYND domain-truncated lines (ZMYND11 Δ/Δ; **Figures 4A and S4b**). Western blot confirmed expression of a truncated ZMYND11 protein lacking the MYND domain (**Figures 4B and S4C**), and the edited lines maintained normal karyotype and pluripotency marker expression (**Figures S4D-S4E**).

**Figure 4.**
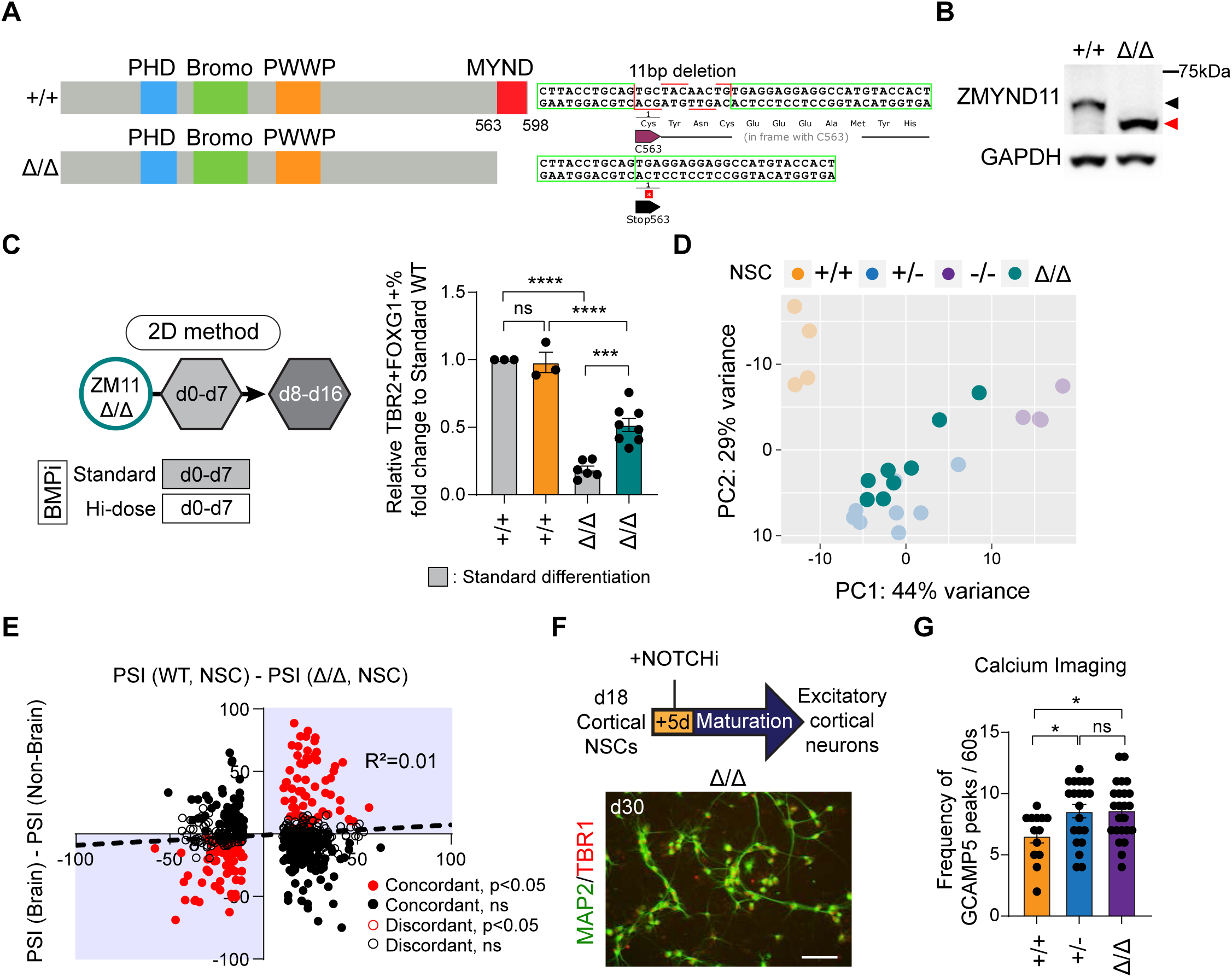
Loss of MYND domain in ZMYND11 impairs neurogenesis and leads to neuronal hyperexcitability. **(A)** Schematic outline of CRISPR editing to introduce premature stop codons at the start of C-terminal MYND domain. **(B)** Western blot on MYND domain truncated *ZMYND11* mutant (ZMYND11 Δ/Δ) vs. wildtype (WT); black arrowhead: wildtype band; red arrowhead: domain deleted band. **(C)** Schematic and quantification of flow cytometry analysis for TBR2 and FOXG1 in monolayer culture at day 16 relative to WT after standard differentiation (mean ± SEM, n=3 for +/+, n=6 for Δ/Δ for standard differentiation; n=3 for +/+, n=8 for Δ/Δ for high-dose BMP inhibition). **(D)** PCA of RNA-seq on ZMYND11 Δ/Δ cortical neural stem cells (NSCs) in comparison with control and ZMYND11 deficient cortical NSCs (n=8 for Δ/Δ). **(E)** Scatter plot of PSI differences between WT vs. Δ/Δ mutant cortical NSCs and brain vs. non-brain tissues. Red/black marks significant (P < 0.05) or non-significant events, respectively. Closed/open circles show matched vs. unmatched mutant and non-brain tissue trends. **(F)** Schematic of excitatory neuron induction from cortical NSCs and immunostaining of MAP2 (green) and TBR1 (red) on excitatory neurons at d30 (n=3). Scale bar, 50μm. **(G)** Calcium imaging by infecting ZMYND11 Δ/Δ with GCAMP5 viruses and quantification of frequency of GCAMP spike per 60 seconds (mean ± SEM, n=13 for +/+, n=21 for +/-, n=24 for Δ/Δ). Each n in **(C)**, **(D)** and **(F)** represents an independent differentiation, and in **(G)** represents an individual neuron across three independent differentiations. *P* values in **(C)** were calculated by one-way ANOVA followed by Sidak’s test, while in (G) by one-way ANOVA followed by Tukey’s test. ns: not significant; \**P*< 0.05; \*\*\**P*< 0.001; \*\*\*\**P*< 0.0001. See also **Figures S4-S5**.

We first asked whether MYND domain loss recapitulates the neurogenesis deficit we previously observed in ZMYND11 mutants. Using intracellular flow cytometry to quantify intermediate progenitor cells (IPCs) production in monolayer cortical differentiations at day 16 (**Figure S4F**), we found that ZMYND11 Δ/Δ lines produced fewer TBR2 positive IPCs than wild-type controls (**Figure 4C**). This deficit was rescued by high-dose BMP inhibition, recapitulating the BMP-dependent rescue we previously demonstrated in heterozygous and knockout mutants (Chang et al., 2025). Transcriptomic analysis of ZMYND11 Δ/Δ cortical NSCs showed de-repression of BMP and WNT signaling (**Figure S5A and Table S1**), and principal component analysis positioned these cells closer to heterozygous mutant NSCs than knockout cells (**Figure 4D**). Together, these results indicate that the MYND domain loss produces a partial loss-of-function state that approximates but does not fully phenocopy complete ZMYND11 deficiency. Notably, alternative splicing analysis of ZMYND11 Δ/Δ cortical neural progenitors did not reveal the non-brain isoform switch observed in knockout neural progenitors (**Figures 4E**, **S5B and Table S2**). Because the MYND domain mediates protein-protein interactions, this result argues against a model in which ZMYND11 regulates tissue-specific splicing primarily through MYND-dependent recruitment of splicing machinery (Lian et al., 2024). Instead, the splicing changes in *ZMYND11* mutants are secondary to altered transcription rather than direct effects on the splicing. We next differentiated ZMYND11 Δ/Δ cortical NSCs toward excitatory neurons to test whether the MYND domain loss is sufficient to drive the hyperexcitability phenotype. These neurons expressed appropriate cortical markers (**Figure 4F**) and exhibited increased spontaneous calcium activity, similar to heterozygous ZMYND11 mutant neurons at day 60 (**Figure 4G**). Transcriptomic and splicing analyses of ZMYND11 Δ/Δ excitatory neurons further confirmed their similarity to heterozygous mutants at both the gene expression (**Figure S5C**) and alternative splicing levels (**Figures S5D-S5E and Table S2**). The convergence between MYND domain loss and heterozygous patient-like mutations across multiple readouts including neurogenesis, transcriptomic profile, and neuronal excitability, suggests that disruption of MYND-mediated interactions accounts for a substantial portion of the phenotypic consequences of ZMYND11 haploinsufficiency in excitatory neurons.

These findings show that the MYND domain is required for normal ZMYND11 functions in neurons and help explain the functional impact of patient missense variants in this region.

## Discussion

Our study demonstrates that *ZMYND11* mutations produce E:I imbalance through selective vulnerability of excitatory cortical neurons rather than parallel disruption of both neuronal subtypes. By systematically comparing excitatory and inhibitory neurons derived from the control and mutant lines, we showed that these two lineages display fundamentally different molecular phenotypes to the same chromatin perturbation whereas diverging not only in the magnitude of transcriptomic disruption but also in the gene programs affected and the directionality of alternative splicing changes. Combined with our previous characterization of ZMYND11 function in cortical neural progenitors, these findings revealed that a single chromatin perturbation can produce different transcriptional and functional outcomes across neuronal lineages.

The observation that ZMYND11 haploinsufficiency substantially disrupts excitatory neuron gene expression and function while leaving inhibitory neurons largely unaffected has implications beyond *ZMYND11* itself. We previously classified chromatin-related autism risk genes into functional categories based on their effects on neurogenesis, and *ZMYND11* groups with *ASXL3* (Woods et al., 2025), *DEAF1* (Nabais Sá et al., 2019), *ASH1L* (Liu et al., 2021), *CUL3* (Blackburn et al., 2025) and *KDM5B* (Borroto et al., 2024) as Class I genes that share progenitor-level phenotypes (Cederquist et al., 2020; Chang et al., 2025). Notably, variants in each of these genes have also been associated with seizures, raising the possibility that cell-type-selective vulnerability in excitatory neurons may be a shared downstream mechanism for this class of chromatin regulators. In support of this idea, a study of the role of neuronal Cul3 in mice found that E:I imbalance with Cul3 haploinsufficiency was initiated by glutamatergic hyperactivity (Dong et al., 2020). Whether this principle is generalizable across Class I chromatin regulators remains to be tested, but our data suggest that deriving both excitatory and inhibitory neurons from the same mutant backgrounds should become standard practice when investigating E:I imbalance in stem cell-based NDD modeling.

Why inhibitory neurons are relatively resistant to ZMYND11 haploinsufficiency is not fully resolved by our data, but several observations provide candidate explanations. Higher baseline *ZMYND11* expression in inhibitory neurons may buffer against heterozygous loss, consistent with the finding that only complete knockout produced clear transcriptomic separation in this lineage. Additionally, the chromatin landscape differs between excitatory and inhibitory neuron lineages, and the availability of H3K36me3, the histone mark recognized by ZMYND11, at specific genomic loci may differ in ways that modulate sensitivity to ZMYND11 dosage. Future studies using CUT&Tag or CUT&RUN for H3K36me3 in both neuronal subtypes could directly test whether differential chromatin context underlies the cell-type selectivity we observed.

Among different domains in ZMYND11, the MYND domain is less well-characterized. Unlike PWWP, bromo, or PHD domains, this domain is not involved in recognition of H3K36me3 (Wen et al., 2014), nor does it interact with DNA (Kateb et al., 2013), even though it has two zinc binding motifs. Several studies have shown that MYND domain interacts with other transcriptional and epigenetic regulators including MGA (Ansieau and Leutz, 2002), NCOR1 (Masselink and Bernards, 2000), HNRNPA1 (Lian et al., 2024), EZH2 (Velasco et al., 2006; Tang et al., 2021), KMT2A (Greben et al., 2025) and NSD1 (Greben et al., 2025), viral proteins including adenoviral oncoprotein E1A (Hateboer et al., 1995) and EBNA2 (Ansieau and Leutz, 2002), as well as *trans-*membrane proteins BMPR1A (Sun et al., 2018). It is interesting to note that MYND domain mediates interaction with both repressor EZH2, mediating H3K27me3 deposition, and activator KMT2A, mediating H3K4me3 deposition. However, EZH2 is potentiated by this interaction (Tang et al., 2021), while KMT2A is repressed by this interaction (Greben et al., 2025). While the underlying mechanism is unclear, this still supports the overall role for ZMYND11 as a transcriptional co-repressor.

The findings that MYND domain loss recapitulated the gene expression in heterozygous mutants but not the splicing phenotype of complete ZMYND11 deficiency are informative in two ways. First, it reinforces the model that alternative splicing changes in ZMYND11 mutants are secondary to transcriptional dysregulation rather than direct effects on splicing machinery, consistent with mass spectrometry data from mouse brain identifying transcriptional regulators rather than splicing factors as the predominant MYND domain interaction partners (Greben et al., 2025). Second, it suggests that patients carrying MYND domain missense mutations may exhibit a subset of the full ZMYND11 loss-of-function phenotype. This has practical relevance for predicting phenotypic severity from genotype, though it remains speculative until patient-derived iPSC models carrying specific MYND domain missense variants are characterized.

Several limitations of our study should also be noted. Our assessment of neuronal excitability relied on GCaMP5-based calcium imaging at a relatively low frame rate (∼0.33-0.67 Hz), which provides a measure of overall spontaneous activity but cannot resolve individual action potentials or distinguish intrinsic excitability from synaptic network effects. Microelectrode array recordings or patch-clamp electrophysiology would enable more precise characterization of firing properties, synaptic currents, and network dynamics in future studies. Additionally, our transcriptomic comparisons between excitatory and inhibitory neurons are complicated by the fact that these populations were derived using different patterning protocols and therefore differ in developmental history, regional identity, and maturation trajectory. While the absence of shared differentially expressed genes between the two lineages is striking, we cannot exclude the possibility that some differences reflect protocol-specific rather than cell-type-intrinsic responses. Co-culture or assembloid models in which excitatory and inhibitory neurons mature within the same environment could help disentangle these variables. Our analysis of alternative splicing directionality relied on tissue-level PSI values from the *VastDB* database (Tapial et al., 2017), which are derived from bulk tissue sequencing and do not distinguish cell type-specific splicing patterns. Because the brain contains substantially more excitatory than inhibitory neurons, *VastDB* "brain" PSI values are biased toward excitatory neuron splicing. The apparent brain-isoform switch in ZMYND11-deficient inhibitory neurons should therefore be interpreted cautiously, and single-cell or long-read sequencing of purified human inhibitory neurons will be needed to establish an appropriate reference for this cell type.

## Resource availability

### Lead contact

Requests for further information and resources should be directed to the lead contact, Jason Tchieu

### Materials availability

All materials used in our analysis is available to any researcher for purposes of reproducing or extending the analysis. Cell lines can be provided through materials transfer agreements (MTAs).

### Data and Code Availability

The RNA sequencing data generated in this paper is uploaded to GEO with accession number GEO: GSE326983. This dataset also includes raw count tables of RNA-seq. This paper does not report original code.

## Acknowledgements

We thank Dr. Kenneth Campbell (CCHMC) and members of the Tchieu lab for their critical reading, helpful critiques, and advice on this study. This work was supported by Cincinnati Children’s Research Foundation (J.T., X.C.) and U.S. Department of Defense grant (W81XWH-22-1-0533 to J.T.).

## Author contributions

Conceptualization: XC, JT; Methodology: XC, WL, IY, MM, EA, EI, FG, AS, JT; Investigation: XC, JT; Visualization: XC, JT; Funding acquisition: JT; Project administration: JT; Supervision: JT; Writing - original draft: XC, JT; Writing - review & editing: XC, WL, IY, MM, EA, EI, FG, AS, JT.

## Methods

### Maintenance and propagation of human embryonic stem cells (hESCs)

Experiments using the human embryonic stem cell (hESC) line H1 (WA01) were performed under protocol EIP190101, approved by the Cincinnati Children’s Hospital Medical Center (CCHMC). The H1 line (NIHhESC-10-0043) was obtained from the WiCell Research Institute. All research was conducted in accordance with institutional guidelines, relevant regulations, and the principles outlined in the 2021 ISSCR Guidelines for Stem Cell Research and Clinical Translation. hESCs were thawed and maintained in Essential 8 (E8, Gibco, A1517001) or Essential 8 flex (Gibco, A2858501) medium on Vitronectin-coated (Gibco, A31804) dishes following established protocols. Briefly, hESCs (H1, XY) were cultured at 37°C in a 5% CO_2_ environment and passaged as clumps in 1:5 ratio using 0.5mM EDTA dissociation solution (Invitrogen, 15575020), with 10 μM Y-27632 dihydrochloride (hellobio, HB2297) for the first 24 hours of culture. Cells were used for differentiation between passage 50-70. All hESC lines are karyotypically normal, tested for mycoplasma every 2-3 months and daily monitored to ensure no bacterial or fungal contamination and no differentiation. Pluripotency checks were regularly done with immunostaining, as described before (Chang et al., 2025). hESCs were frozen using E8 medium + 10% DMSO. *ZMYND11* genotyping by Sanger sequencing was performed regularly to verify the cell identity.

### Human embryonic kidney 293T (HEK293T) cell culture and lentivirus production

HEK293T cells were cultured at 37°C in a 5% CO_2_ environment using DMEM medium (Gibco, 11960044) supplemented with 10% fetal bovine serum (FBS, R&D Systems, S11150), 1X Antibiotic-Antimycotic (Gibco, 15240062) and 1X GlutaMAX (Gibco, 35050061). The medium was changed every 2 days, and cells were passaged using Trypsin-0.05% EDTA (Gibco, 25300054).

For lentivirus production, the day before transfection, 5 million HEK293T cells were plated in a 10 cm dish coated with Poly-L-lysine (Sigma-Aldrich, P4832) & 0.1% gelatin (Sigma-Aldrich, G1890). The hSYN1-GCAMP5-P2A-mCherry, packaging (psPAX2) and envelope (pMD2.G) vectors were transfected using X-tremeGene 9 (Sigma-Aldrich, 6365779001) in a 1:2:1 molar ratio, respectively. Medium was changed into 20 μg/ml BDNF (R&D Systems, 248-BDB), 20 μg/ml GDNF (PeproTech, 450-10) and 200 μM Ascorbic acid (Sigma-Aldrich, A8960) in Neurobasal medium (Gibco, 21103049) supplemented with B27 (1:50, without Vitamin A, Gibco, 12587010; NB-B27) and 1X GlutaMAX 18 hours post transfection. Viruses were harvested at 48 and 72 hours post transfection, concentrated using AMICON Ultra-15 Centrifugal Filter Units (Millipore, UFC9100), and snapped frozen in liquid nitrogen.

### Monolayer prefrontal cortical (PFC) differentiation, excitatory neuron induction and inhibitory neuron differentiation

Dishes were coated with Matrigel (Corning, 354234, diluted 1:100 in DMEM/F12, Gibco, 11330032) and stored overnight at 4°C. The next day, hESCs were dissociated to single cells using Accutase (Sigma-Aldrich, A6964), resuspended with E8 and plated on the Matrigel coated dish at a density of 300,000 cells /cm^2^ in the presence of 10 μM ROCKi on day -1. From day 0 to 2, the cells were cultured in Essential 6 (E6, Gibco, A1516401) medium supplemented with 100 nM LDN193189 (R&D Systems, 6053), 10 μM SB431542 (hellobio, HB3555) and 5 μM XAV939 (hellobio, HB0660) (i.e., LSBX) to promote anterior telencephalic patterning. From day 3 to 7, the cells were maintained in E6 with LSB with XAV939 removed. From day 8 to 18, the cells were treated with 50 ng/ml FGF8 (PeproTech, 100-25) in N2 medium supplemented with B27 (1:1000, without Vitamin A; N2-B27) to promote a PFC-like cortical regionalization.

For hi-dose BMP inhibition, 500 nM LDN193189 was used from day 0 to 7 during differentiation.

To induce cortical neural stem cells (NSCs) to neurons, 10 μM DAPT (NOTCHi, hellobio, HB3345) was added to day 18 of hi-dose BMP inhibition differentiation in NB-B27 for 5 days. The neurons were then replated onto dishes coated with Poly-ornithine (PO, Sigma-Aldrich, P3655), Laminin (Lam, R&D Systems, 3400-010-02) and Fibronectin (FN, R&D Systems, 1918-FN) and maintained in NB-B27 supplemented with 20 μg/ml BDNF, 20 μg/ml GDNF and 200 μM Ascorbic acid. To eliminate proliferative cells from the neuron culture, 1 μM Ara-C (Sigma-Aldrich, C6645) was added for 48 hours. Fifty percent of neuron medium was changed every 3 days.

For inhibitory neuron differentiation, 1uM sonic hedgehog agonist (SAG, bio-techne, 4366) was added to the cortical differentiation from day 8 to 18. Neuronal induction was similarly performed as excitatory neurons.

### Calcium imaging of *ZMYND11* mutant neurons and analysis

hPSC-derived excitatory neurons were infected with hSYN1-GCAMP5-P2A-mCherry lentiviruses and cultured on 8-well μ-slide plate (ibidi, 80826). For each batch of experiments, both mutant and control excitatory neurons were seeded at equal number and infected with viruses at the same time to account for the variability in the expression of GCAMP5. Imaging of these neurons was performed after mCherry expression was confirmed, indicating these neurons expressed synapsin-1 and were relatively mature (d60+). To record spontaneous activities, GCAMP5 fluorescence was recorded under fluorescence microscope at a frame rate of around 1 frame per 3 second in Fig. 1d and 1.5 second in Fig. 1f. The time-course data was analyzed in ImageJ and region of interest was placed on neuronal soma to calculate GCAMP5 activities during the time course. To quantify the percentage of firing neurons, the number of neurons that showed spontaneous calcium handling (GCAMP5 spiking) was divided by the number of neurons that were mCherry positive. To quantify firing frequency, the signal intensity calculated in each frame was normalized to the baseline fluorescence levels to detect spikes, which was visualized by line plots.

### Generation of *ZMYND11* mutant hESCs

*ZMYND11* heterozygous mutant and knockout lines were as previously described (Chang et al., 2025). For MYND domain truncated *ZMYND11* mutant, CRISPR/Cas9 was used to introduce frameshift mutations into last exon of ZMYND11 that encodes MYND domain in H1 hESCs. Guide RNA (gRNA, IDT): 5’-CCTGCAGTGCTACAACTGTG was chosen based on its location near the start of the exon. The gRNA-Cas9 (IDT, 1081060) complex was transfected using Lipofectamine Stem Transfection Reagent (Invitrogen, STEM00003) according to manufacturer’s protocol. Individual clones were isolated by replating transfected cells at low density. Indels were screened using Sanger sequencing. Selected clones were clonally expanded and differentiated using above-mentioned protocol.

### Histology and Immunohistochemistry

Cells were replated onto glass coverslip 24 hours before fixation. The cells were fixed in 4% PFA (Fisher Scientific, 50-980-487) for 10 minutes at 4°C, permeabilized with 0.5% Triton X-100 (Fisher Scientific, BP151) for 5 minutes and blocked with 5% NDS (Sigma-Aldrich, S30-100ML) in 0.2% Tween-20 (Fisher Scientific, BP337) for 1 hour. The cells were then incubated with primary antibodies overnight at 4°C. The cells were then washed 3 times with 0.2% Tween-20 and incubated with secondary antibodies for 30 minutes at room temperature, washed and stained with DAPI (Sigma-Aldrich, D9542). The cells were then embedded in Fluoromount (SouthernBiotech, 0100-01) and stored at 4°C until imaging.

### Western blot

Cells were dissociated and counted. An equal number of cells were harvested per experiment, lysed in 1X LDS Sample Buffer (Invitrogen, B0007) supplemented with 1X Sample Reducing Agent (Invitrogen, B0007) then denatured at 95°C for 5 minutes. Samples were then loaded and run on a 4-12% pre-cast Bis-Tris gel (Invitrogen, NW04125BOX) and transferred onto Nitrocellulose membrane (Fisher Scientific, 15255928) for 2 hours. Membranes were blocked in 5% milk (US Biological, S1013-90A) and primary antibodies were incubated overnight at 4°C. The next day membranes were washed with 1X TBS buffer (BIO-RAD, 1706435) with 0.1% Tween-20 and incubated with HRP-conjugated secondary antibodies for 1 hour. Blots were visualized using SuperSignal Chemiluminescent Substrate (Thermo Scientific, 34580).

### Intracellular FACS analysis

Cells were dissociated using Accutase and washed with DPBS (Gibco, 14190235). Approximately 5 x 10^5^ cells were treated with Zombie Violet Live/Dead Dye (1:1000, Biolegend, 423113) in DPBS for 30 minutes. After washing with DPBS, the cells were fixed using 1X Fix/Perm buffer from the BD Pharmingen Transcription Factor Buffer Set (BD, 562574) and incubated at 4°C for 1 hour. Cells were then washed twice with 1X Perm/Wash buffer and incubated with primary antibodies for 1 hour. After 2 more washes, secondary antibodies were added, and cells were incubated at 4°C for 1 hour in the dark. Finally, the cells were washed 2 times and prepared for analysis.

### RNA extraction, RNA-seq and gene expression analysis

RNA was extracted using TRIzol (Invitrogen, 15596018) followed by chloroform (Sigma-Aldrich, C2432) extraction. Purification of RNA was performed using RNeasy Kits (Qiagen, 74104) and eluted in nuclease free water. Total RNA was submitted to Novogene for paired-end sequencing (poly-A enriched, 150bp). Raw FASTQ files were quality checked using FastQC. Adapter sequences were trimmed using fastp with parameters “-q 20 -n 7”. Trimmed reads were then aligned to ENSEMBL GRCh38 using HISAT2 with default parameters. A raw count file was generated using featureCounts in Subread with default parameters and genes with no raw counts in >= 75% of samples were filtered out. Downstream analysis was done using R packages. Normalized counts per million (cpm) values were generated using “cpm” in “edgeR”. Principal component analysis was performed using “plotPCA” in “DESeq2”. Differential gene expression analysis was performed using “DESeq2” with *P*.adjust < 0.05 and fold change > 1.5. Gene ontology analysis was performed using “clusterProfiler” with *P*.adjust calculated using Benjamini-Hochberg method. Plots were generated using “ggplot2”.

Reanalysis of *ZMYND11* mutant and control ESCs and NSCs was based on deposited datasets in GEO (GSE279072).

### Alternative splicing analysis using rMATS

Input alignment files for rMATS splicing analysis were generated either using HISAT2 with trimmed reads or STAR with raw reads with default parameters. Results from rMATS were further filtered by removing low coverage events (avg_read >5) using “maser” in R. Significant splicing events were identified with FDR < 0.05 and percent spliced-in change (ΔPSI) > 0.1.

Sashimi plots were generated using rmats2sashimiplot. For exon usage principal component analysis, PSI matrix was generated using rMATS with parameters “--statoff”. PSI matrix was then imputed by KNN method for missing values using “impute” in R. PCA was conducted using “prcomp” function in R.

### Alternative splicing analysis using AltAnalyze

For AltAnalyze we only input BAM files generated by STAR as indicated by user manual. Significant splicing events were identified with raw *P* value < 0.05 and ΔPSI > 0.1. Events from AltAnalyze were more than those from rMATS as AltAnalyze summarized events at each individual exon-exon or exon-intron junction while rMATS summarized events of individual exon. Therefore, a significant changed spliced exon or intron might have more than one event reported in AltAnalyze.

Overlapping significant spliced exon events by STAR-AltAnalyze with those from HISAT2-rMATS or STAR-rMATS was performed using genomic coordinates of the target exon. To expand the potential overlap, events that have matched upstream & downstream exon coordinates and either matched 5’ target exon coordinate or 3’ target exon coordinate were also included. Plus, events predicted from different methods would have to be in the same direction (both predicted to be spliced-in or spliced-out) to be considered as overlapped. These overlapped events were subsequently called high confidence differential alternative splicing (DAS) events for further analysis.

Tissue-specific PSI values for high confidence DAS events were extracted from *VastDB* database (https://vastdb.crg.eu) according to target exon coordinates in ENSEMBL GRCh38 and categorized into either brain tissue or non-brain tissue for comparison. Heatmap was generated using “pheatmap” in R (row-wise scaled by Z-score transformation) after missing values were imputed by KNN method using “impute” in R. Correlation plots were generated using Prism.

### Quantification, statistical analysis and reproducibility

All data represent mean ± SEM. Statistical significance was determined by parametric tests including unpaired two-tailed t test for two groups and analysis of variance (ANOVA) for more than two groups with post hoc tests as indicated. A *P* value <0.05 was considered statistically significant. *P.*adjust values in gene ontology analysis were calculated using Benjamini-Hochberg method. Prism 8 software was used for statistical analysis. For all cell culture experiments, n represents total number of independent differentiations. Each experiment was repeated at least 3 times.

### Supplemental information

Document S1. Figures S1-S5

Table S1. Differential gene expression analysis.

Table S2. Alternative splicing analysis.

Table S3. Primary and secondary antibodies used in this study.

Video S1. Calcium imaging in WT excitatory neurons at d60.

Video S2. Calcium imaging in *ZMYND11* +/− excitatory neurons at d60.

Video S3. Calcium imaging in *ZMYND11* −/− excitatory neurons at d60.

Video S4. Calcium imaging in WT excitatory neurons at d120.

Video S5. Calcium imaging in *ZMYND11* +/− excitatory neurons at d120.

---

**Figure S1.**
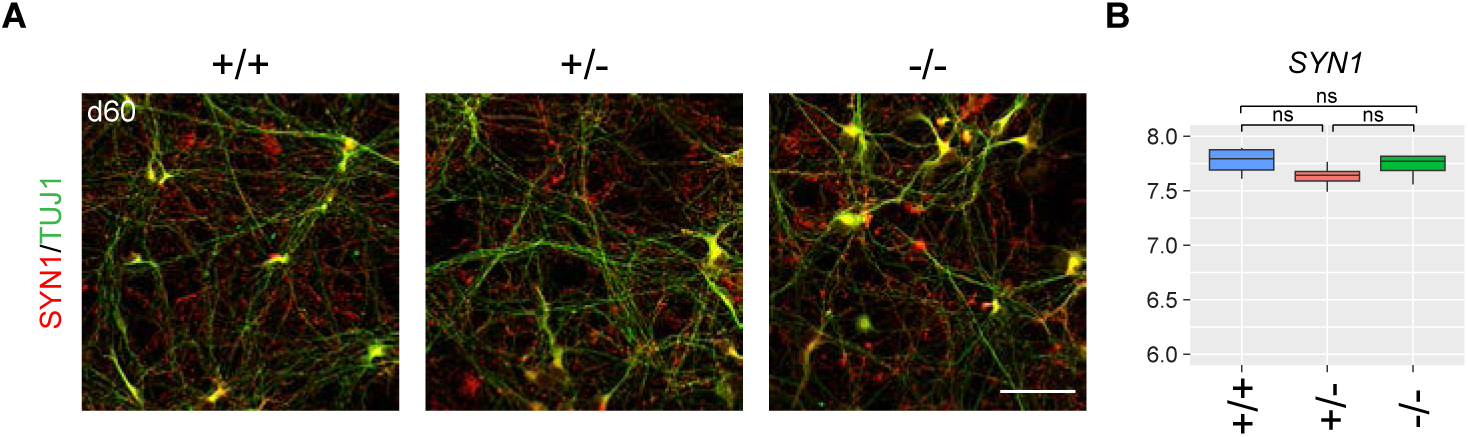
ZMYND11 deficient excitatory neurons show similar maturation with control. **(A)** Immunostaining of TUJ1 (green) and SYN1 (red) on excitatory neurons at day 60 (d60, n=3). Scale bar, 50μm. **(B)** Normalized expression of *SYN1* in excitatory neurons from transcriptome. Boxplot shows median, quartiles and whiskers (min/max) values. Each n represents an independent differentiation. *P* values in **(B)** were calculated by DESeq2 statistic test. ns: not significant.

**Figure S2.**
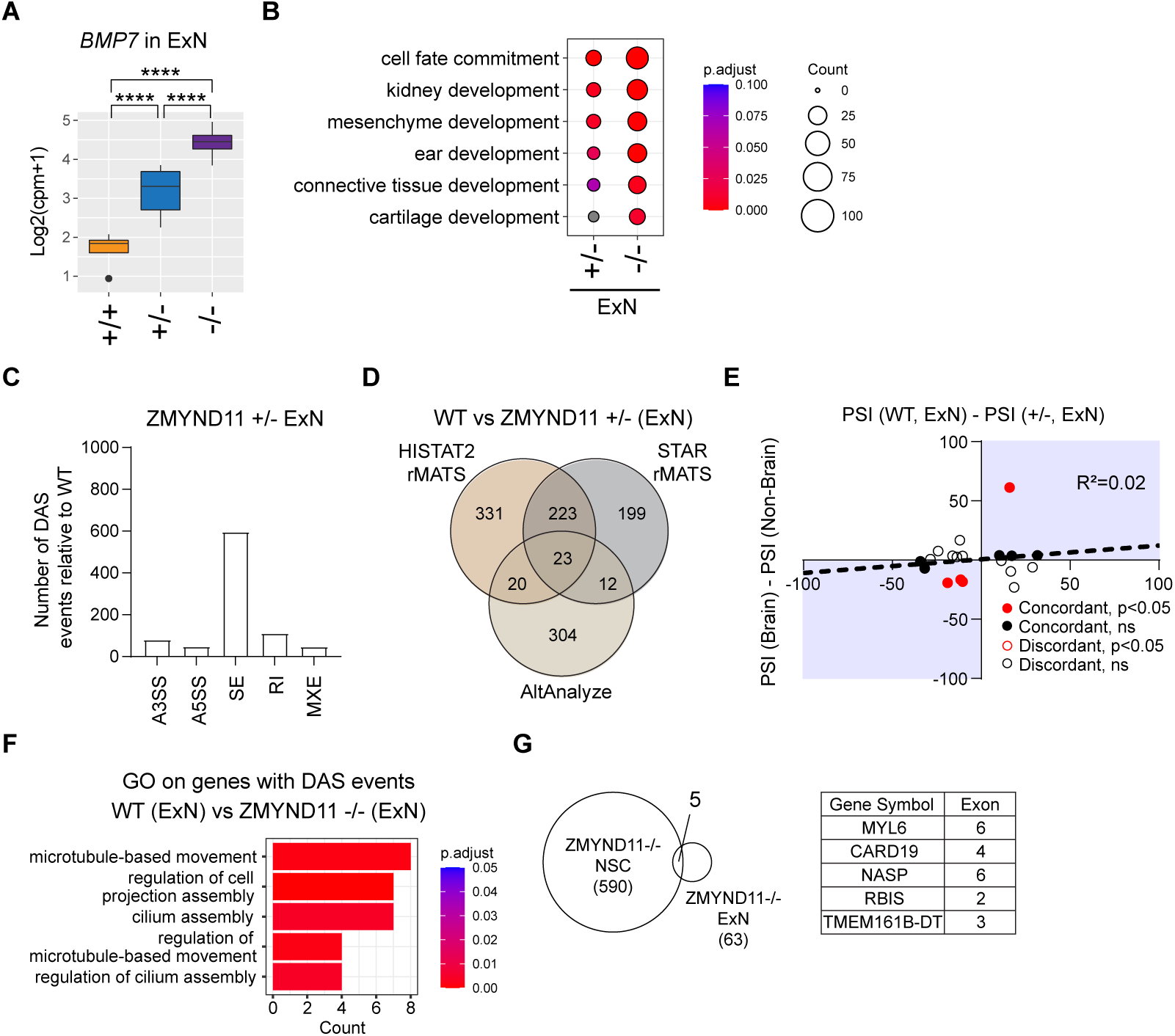
Additional transcriptomic analysis on ZMYND11 deficient excitatory neurons. **(A)** Normalized expression of *BMP7* that are upregulated in ZMYND11 deficient excitatory neurons. Boxplot shows median, quartiles and whiskers (min/max) values. **(B)** Gene ontology analysis on ZMYND11 deficient excitatory neurons focusing on non-brain development terms. Gray dots: *P*.adjust>0.1. **(C)** Number of differential alternative splicing (DAS) events in ZMYND11 +/- excitatory neurons (rMATS-HISAT2). **(D)** Overlap of DAS events predicted by HISAT2-rMATS, STAR-rMATS and AltAnalyze in ZMYND11 +/- excitatory neurons. **(E)** Scatter plot of PSI differences between WT vs. heterozygous mutant excitatory neurons and brain vs. non-brain tissues. Red/black marks significant (P < 0.05) or non-significant events, respectively. Closed/open circles show matched vs. unmatched mutant and non-brain tissue trends. **(F)** Gene ontology analysis on genes with high-confidence DAS events in ZMYND11 -/-excitatory neurons. **(G)**. Overlap of high-confidence DAS events between ZMYND11 -/- cortical NSCs and excitatory neurons. *P* values in **(A)** were calculated by DESeq2 statistic test. \*\*\*\**P*<0.0001.

**Figure S3.**
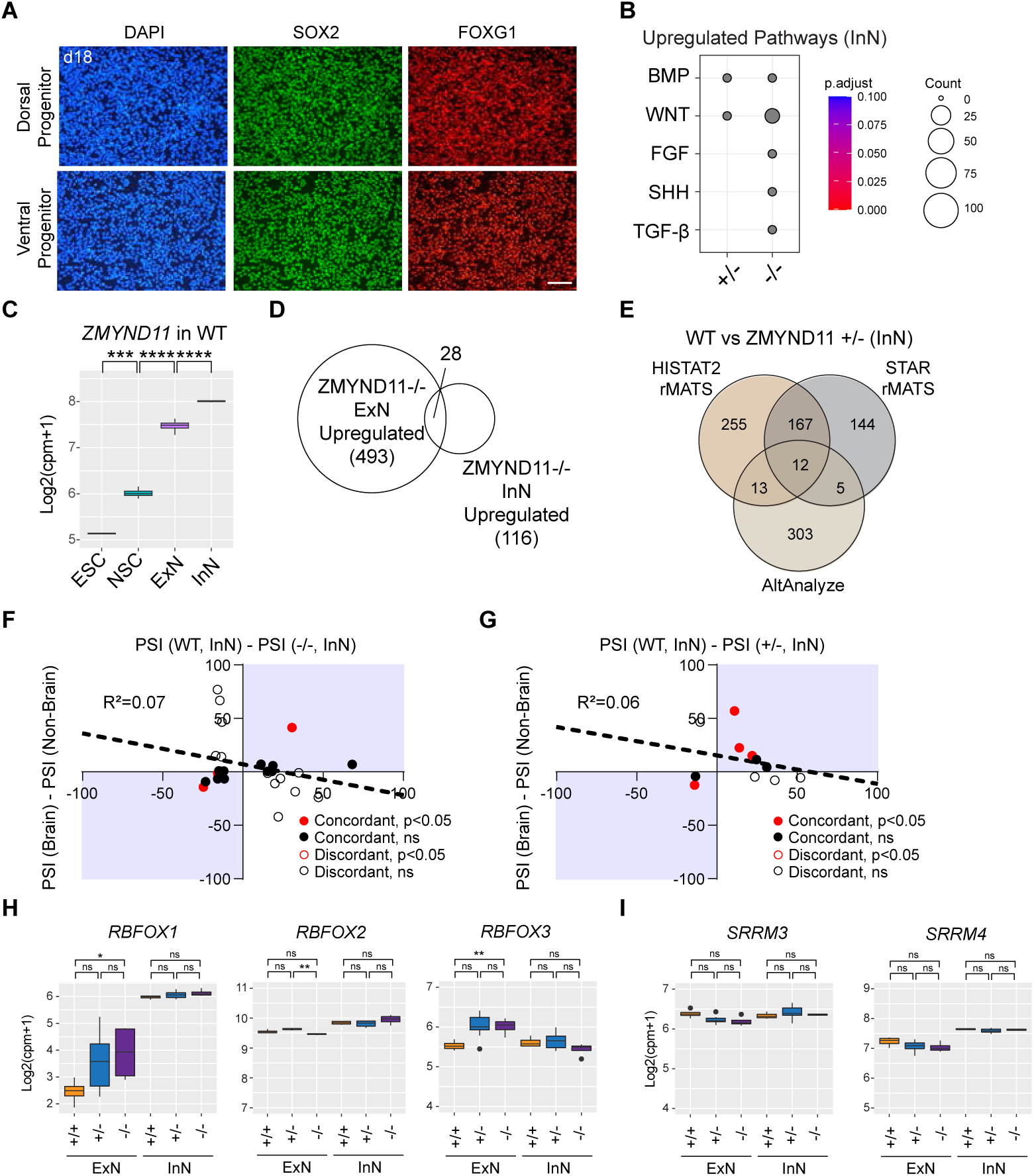
Additional transcriptomic analysis of ZMYND11 deficient inhibitory neurons. **(A)** Immunostaining of SOX2 (green) and FOXG1 (red) on wildtype (WT) dorsal progenitors and ventral progenitors at day 18 (d18, n=3, Each n represents an independent differentiation). Scale bar, 50μm. **(B)** Gene ontology analysis on upregulated genes in ZMYND11 deficient inhibitory neurons. Gray dots: *P*.adjust > 0.1. **(C)** Normalized expression of *ZMYND11* in WT cells. Boxplot shows median, quartiles and whiskers (min/max) values. **(D)** Overlap of upregulated genes between ZMYND11 -/- excitatory neurons and inhibitory neurons. **(E)** Overlap of differential alternative splicing (DAS) events predicted by HISAT2-rMATS, STAR-rMATS and AltAnalyze in ZMYND11 +/- inhibitory neurons. **(F-G)** Scatter plot of PSI differences between WT vs. knockout mutant **(F)** or heterozygous mutant **(G)** inhibitory neurons and brain vs. non-brain tissues. Red/black marks significant (P < 0.05) or non-significant events, respectively. Closed/open circles show matched vs. unmatched mutant and non-brain tissue trends. **(H)** Normalized expressions of *RBFOX1*, *RBFOX2* and *RBFOX3* in neurons. Boxplot shows median, quartiles and whiskers (min/max) values. **(I)** Normalized expressions of *SRRM3* and *SRRM4* in neurons. Boxplot shows median, quartiles and whiskers (min/max) values. *P* values in **(C)**, **(H)**, **(I)** were calculated by DESeq2 statistic test. \**P*< 0.05; \*\**P*< 0.01; \*\*\**P*< 0.001; \*\*\*\**P*< 0.0001.

**Figure S4.**
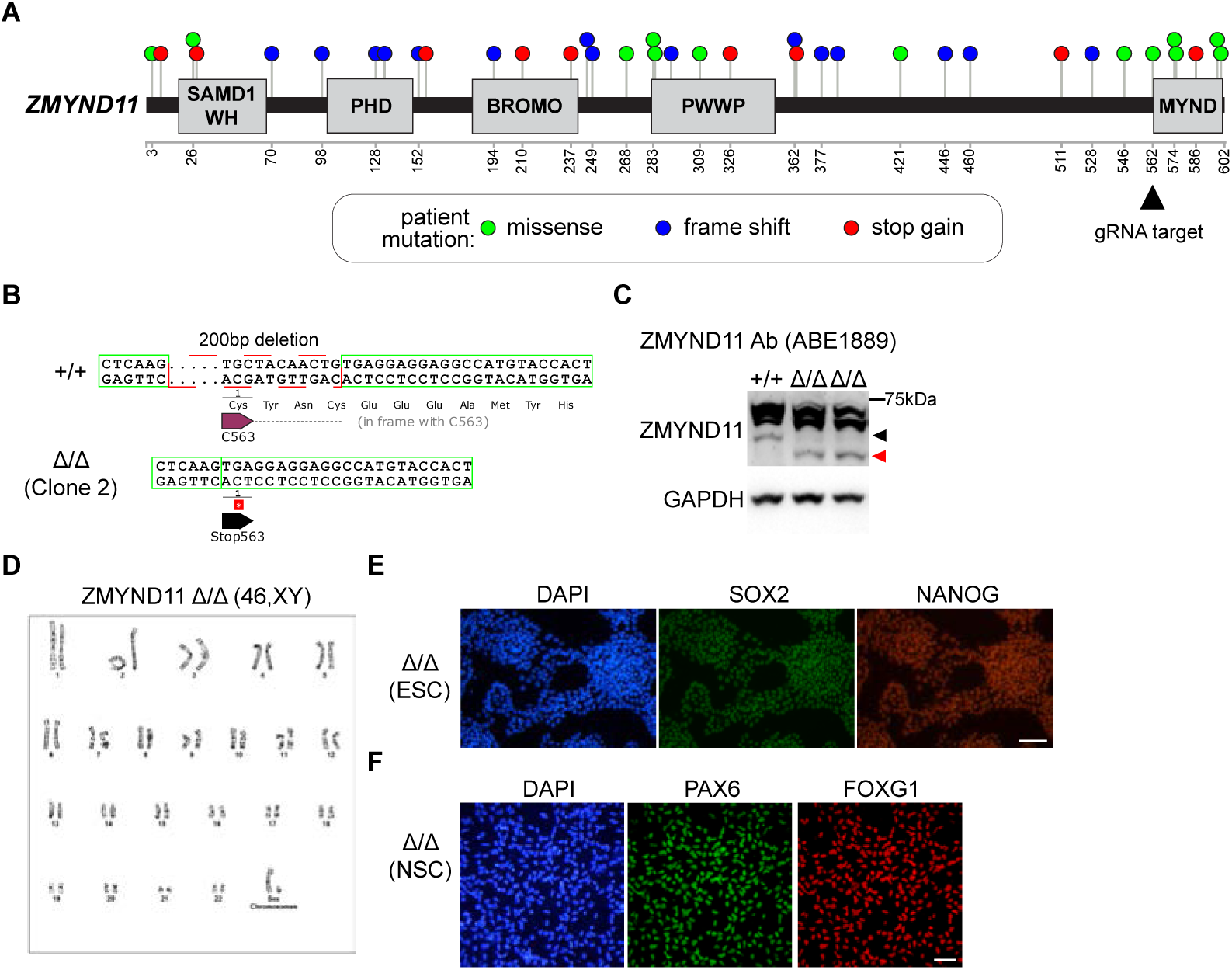
Characterization of ZMYND11 Δ/Δ mutant. **(A)** Patient ZMYND11 mutations distribution from 3 SFARI database (https://gene.sfari.org/, data extracted in May 2024). Black arrowhead marks the location of gRNA targeting the start of MYND domain. Modified from Chang et al., 2025. **(B)** Genotype of the second clone of MYND domain truncated *ZMYND11* mutant. **(C)** Western blot on MYND domain truncated *ZMYND11* mutant (ZMYND11 Δ/Δ) vs. wildtype (WT) using a second ZMYND11 antibody; black arrowhead: wildtype band; red arrowhead: domain deleted band. **(D)** Karyotype of ZMYND11 Δ/Δ mutants. **(E)** Immunostaining of SOX2 (green) and NANOG (red) on ZMYND11 Δ/Δ ESCs (n=3). Scale bar, 50μm. **(F)** Immunostaining of PAX6 (green) and FOXG1 (red) on ZMYND11 Δ/Δ cortical neural stem cells NSCs (n=3). Scale bar, 50μm. Each n represents an independent differentiation.

**Figure S5.**
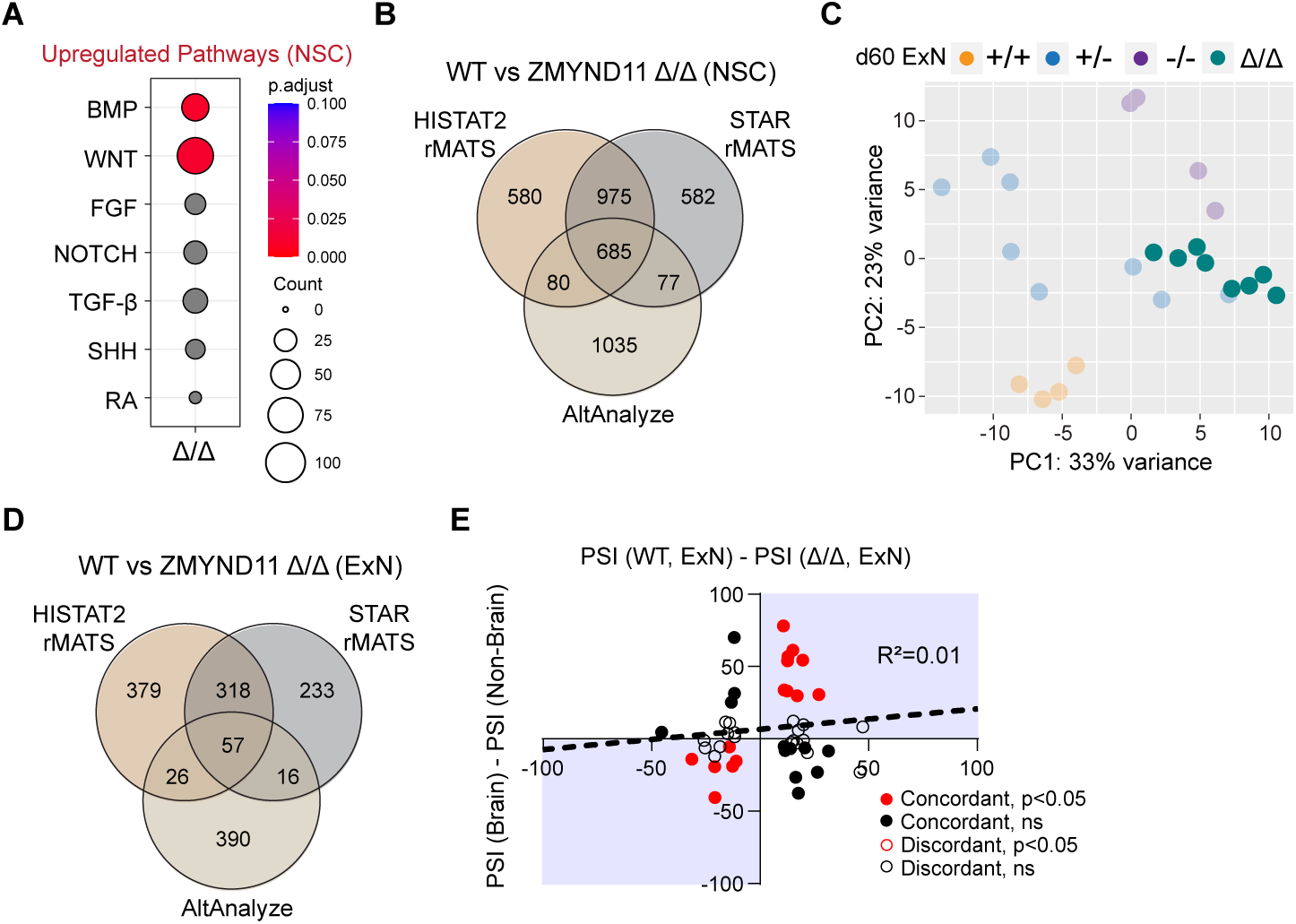
Additional transcriptomic analysis of ZMYND11 Δ/Δ mutant. **(A)** Gene ontology analysis on upregulated genes in ZMYND11 Δ/Δ cortical neural stem cells (NSCs). Gray dots: *P*.adjust > 0.1. **(B)** Overlap of differential alternative splicing (DAS) events predicted by HISAT2-rMATS, STAR-rMATS and AltAnalyze in ZMYND11 Δ/Δ NSCs. **(C)** PCA of RNA-seq on ZMYND11 Δ/Δ excitatory neurons in comparison with control and *ZMYND11* deficient excitatory neurons at day 60 (d60, n=8 for Δ/Δ, Each n represents an independent differentiation). **(D)** Overlap of DAS events predicted by HISAT2-rMATS, STAR-rMATS and AltAnalyze in ZMYND11 Δ/Δ excitatory neurons. **(E)** Scatter plot of PSI differences between WT vs. Δ/Δ mutant excitatory neurons and brain vs. non-brain tissues. Red/black marks significant (*P*< 0.05) or non-significant events, respectively. Closed/open circles show matched vs. unmatched mutant and non-brain tissue trends.

